# Aggregation of Disrupted in Schizophrenia 1 arises from a central region of the protein

**DOI:** 10.1101/2023.08.03.551216

**Authors:** Beti Zaharija, Nicholas J. Bradshaw

## Abstract

An emerging approach to studying major mental illness is through proteostasis, with the identification of several proteins that form insoluble aggregates in the brains of patients. One of these is Disrupted in Schizophrenia 1 (DISC1), a neurodevelopmentally-important scaffold protein, and the product of a classic schizophrenia risk gene. DISC1 was seen to aggregate in post mortem tissue from patients with schizophrenia, bipolar disorder and major depressive disorder, as well as in a variety of model systems, although the mechanism by which it does so is still unclear. Aggregation of two other proteins implicated in mental illness, TRIOBP-1 and NPAS3, was shown to be dependent on very specific structural regions of the protein. We therefore looked to the recently determined domain structure of DISC1, and investigated which structural elements were key for its aggregation. While none of the known DISC1 regions (named D, I, S and C respectively) formed aggregates individually when expressed in neuroblastoma cells, the combination of the D and I regions, plus the linker region between them, formed visible aggregates. Further refinement revealed that a region of approximately 30 amino acids between these two regions is critical to aggregation, with deletion of this region from full length DISC1 sufficient to abolish its aggregation propensity. This finding from mammalian cell culture contrasts with the recent determination that the extreme C-terminal of DISC1 can aggregate *in vitro*, although we did see some indication that combinations of C-terminal DISC1 regions can also aggregate in our system. It therefore appears likely that DISC1 aggregation, implicated in mental illness, can occur through at least two distinct mechanisms.

## Introduction

An emerging theme in the study of major mental illness, is that chronic instances of these conditions may be associated with disrupted proteostasis in the brain [1]. Post mortem analysis of brain tissue from patients with schizophrenia has demonstrated a subgroup of them to have higher levels of insoluble protein [2], suggestive of protein misfolding and aggregation. In support of this, other studies have shown such patients to have poorer proteasome function, and increased levels of ubiquitinated protein, suggestive of clearance of misfolded protein [3,4]. More specifically, several proteins have been identified as being insoluble, and therefore likely aggregating, in the brains of patients with schizophrenia, bipolar disorder and/or major depressive disorder, although each protein appears in only a subset of patients [5–12]. This suggests a scenario similar to neurodegenerative disorders such as amyotrophic lateral sclerosis or frontotemporal lobe dementia, in which a number of different proteins aggregate in the brains of different subgroups of patients. One major difference from this, however, is that mental illnesses are not characterised by largescale neuronal death, meaning that any proteins aggregating in these conditions cannot be neurotoxic in the same manner as those seen in neurodegenerative disorders [1].

The first protein implicated as aggregating in mental illness was Disrupted in Schizophrenia 1 (DISC1), the product of a gene originally found to be destroyed by a chromosomal translocation in a family with high instances of various major mental illnesses [13], and which has since been implicated by genetic evidence in specific populations [14,15]. DISC1 is a scaffold protein that binds to and regulates the expression of many other proteins, a large number of which are involved in neuronal development and/or signalling [16,17]. Given that the *DISC1* gene is implicated in mental illness, the DISC1 protein was selected for an initial study looking at protein aggregates in mental illness [5], which revealed DISC1 to exist in an insoluble state in the brains of a subset of patients with schizophrenia, bipolar disorder and major depressive disorder, but not control individuals [5]. This insolubility corresponds to the formation of visible aggregates, both in *in vitro* models and in a transgenic rat model [6,7,18], with the rat displaying a range of behavioural phenotypes consistent with disrupted dopamine signalling [18–23].

The *DISC1* gene is subject to extensive alternate splicing [24], however the longest commonly studied splice variant, sometimes referred to as the L variant, encodes an 854-amino acid protein [13] that forms higher order oligomers [25,26]. This protein, which will be referred to as full length DISC1, is predicted to be largely disordered in its N-terminal third, with the remainder of the protein forming coiled-coils [13,27,28]. Experimental evidence for the structure of DISC1 is sparse, beyond some NMR structures from the extreme C-terminal end of the protein, showing its interaction with protein binding partners [29,30]. The overall domain structure of DISC1 has been investigated, however, using ESPRIT (Expression of Soluble Proteins by Random Incremental Truncation), in which thousands of truncated versions of a protein are expressed in *E. coli*, and screened for those that form soluble proteins, on the grounds that compactly folded protein domains are more likely to form stable, soluble recombinant proteins, as opposed to partially formed domains [31]. This approach identified four seemingly stable folded domains or regions within human DISC1, labelled D, I, S and C, of which D lies near the centre of the protein, and the other three exist in the C-terminal half of the protein (figure 1) [32].

**Figure 1:**
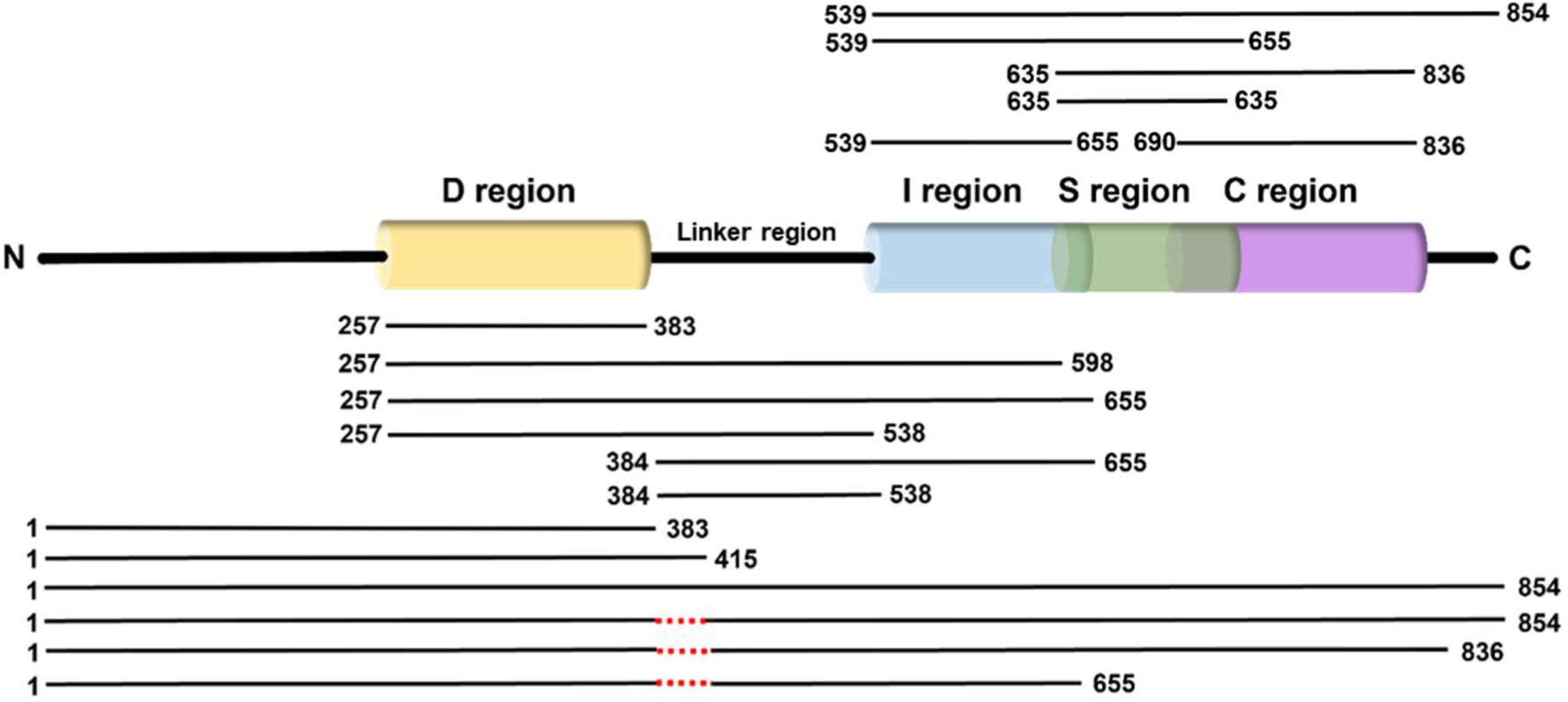
Schematic representation of the one-dimensional structure of DISC1, featuring the four structured regions (D, I, S and C) identified by Yerabham *et al* [32]. A linker region between the D and I regions is also labelled. Black horizontal lines represent the regions of DISC1 encoded for by the various plasmid deletion constructs used in this work.

Notably, for two other proteins known to aggregate in mental illness, TRIOBP-1 and NPAS3, it has been possible to identify specific regions of the proteins that are required for aggregation, with deletion of these regions abolishing aggregation [9–11,33]. It is therefore likely that one or more regions of DISC1 are also responsible for its aggregation propensity. We therefore set out to test, within the context of the proposed domain structure for DISC1, which structural region(s) of DISC1 are likely to account for its aggregation propensity in the brain.

## Materials & Methods

### Plasmids

A Gateway entry vector encoding human DISC1 came from the DNASU Plasmid Repository (Tempe, AZ, USA, clone HsCD00516321 [34,35]), and a plasmid for expressing full length DISC1 in mammalian cells has been described previously [12]. Fragments of *DISC1* were subcloned from existing vectors into pENTR1A no ccDB [36] (Dr. Eric Campeau, via Addgene, Watertown, MA, USA, clone 17398) at the *SalI* and *XbaI* sites, according to a previously described strategy [37]. Entry vectors were transferred using LR clonase recombination (Thermo Fisher Scientific, Walton, MA, USA) into the vectors pdcDNA-FlagMyc (B. Janssens, BCCM/LMBP Plasmid Collection clone LMBP 4705, Zwijnaarde, Belgium) and/or pETG10A (A. Geerlof, EMBL, Heidelberg, Germany). A complete list of plasmids and primers used in this study can be found in supplementary tables S1 and S2, and the regions of DISC1 covered are shown schematically in figure 1. All plasmids were confirmed by sequencing.

### Antibodies

The anti-Flag primary antibody was purchased from Merck (Darmstadt, Germany: Flag F3165). Secondary antibodies were purchased from Thermo Fisher Scientific.

### Cell culture

Human HEK293 kidney cells were grown in D-MEM/10% FCS, supplemented with penicillin and streptomycin. Human SH-SY5Y neuroblastoma cells were grown in D-MEM/F-12/10% FCS, supplemented with L-glutamine, non-essential amino acids, penicillin and streptomycin. All media and supplements were purchased from PAN-Biotech (Aidenbach, Germany) or Thermo Fisher Scientific. Cells were transfected with plasmids using either Metafectene or Metafectene Pro (Biontex, Munich, Germany), according to manufacturer’s instructions.

### Immunocytochemistry & microscopy

Cells on glass coverslips were fixed for 15 minutes with PBS / 4% paraformaldehyde, and permeabilised for 10 minutes with PBS / 1% Triton X-100. Cells were then blocked for 50 minutes with PBS / 10% goat serum, before being stained with the primary antibody diluted in PBS/10% goat serum, and incubated for three and a half hours. Following incubation, cells were washed three times (5 minutes per wash) with PBS. Cells were then stained with DAPI, Acti-Stain 488 (Cytoskeleton, Denver, CO, USA) and/or secondary antibodies in PBS / 10% goat serum for 1 hour, before 3 additional washes with PBS. Coverslips were mounted using Fluoroshield mounting medium (Merck), and viewed on an IX83 Inverted Microscope (Olympus, Shinjuku, Japan) using an ORCA-R2 CCD camera (Hamamatsu Photonics, Hamamastu, Japan) and cellSens imaging software (Olympus).

### Cell lysis

Cells were lysed for 1 hour in PBS / 1% Triton X-100 / 20 mM magnesium chloride, containing DNaseI (New England Biolabs, Ipswich, MA, USA) and protease inhibitor cocktail (Thermo Fisher Scientific), and then denatured by mixing 1:1 with 156 mM Tris pH 6.8 / 5% SDS / 25% glycerol / 100 mM DTT / bromophenol blue and heating to 95 °C for 5 minutes.

### Insoluble protein purification assay

Protein aggregation in transfected HEK293 cells was investigated by lysing the cells and then subjecting them to a set of ultracentrifugation steps, in which soluble protein in the supernatant was separated from insoluble protein in the pellet. Various buffers are used to solubilise the protein between centrifugations, and the full protocol has been published previously [10].

### SDS-PAGE and Western blot

Protein samples were run on bis-acrylamide gels and then transferred to PVDF membranes (Macherey-Nagel, Düren, Germany) using a Trans-blot Turbo Transfer System (Bio-Rad, Hercules, CA, USA). Membranes were blocked in PBS / 0.05% Tween-20 / 5% milk powder for one hour, and stained overnight with primary antibodies in PBS / 0.05% Tween-20. Membranes were then washed 3 times over 30 minutes in the same buffer, before being stained with secondary antibodies and washed 3 more times, again using the same buffer. Protein was visualised with ECL (Thermo Fisher Scientific) on a ChemiDoc MP Imaging System, using Image Lab software (Bio-Rad), which was also used for quantification where needed.

## Results

As expected, the full length DISC1 protein showed clear cytoplasmic aggregates when expressed in SH-SY5Y neuroblastoma cells (figure 2A-B). In contrast, none of the four structural domains previously predicted by ESPRIT signalling showed any indication of protein aggregation when expressed under the same conditions (D: amino acids, AA, 257-383, I: AA 539-655, S: AA 635-738, C: AA 691-836, figure 2C-G), indicating that none of these alone is capable of inducing aggregation, at least in this system.

**Figure 2:**
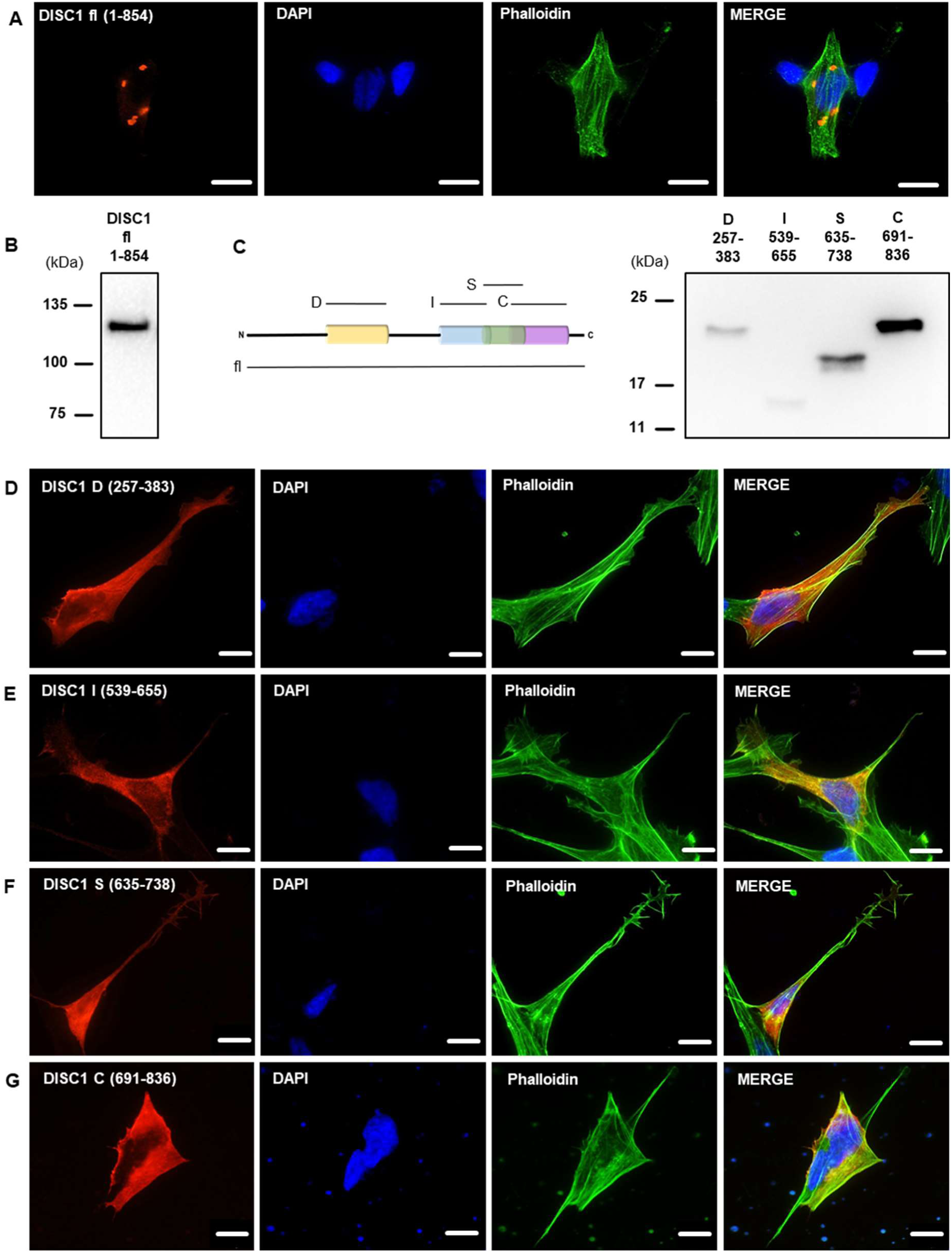
No individual structured domain of DISC1 aggregates in isolation. (**A**) Full length DISC1 forms aggregates in the cell. (**B**) Western blot of the full length DISC1 construct. (**C**) Constructs encoding individual regions of DISC1, shown in schematic form and by Western blot. (**D**) The D region does not readily aggregate when expressed alone. (**E**) The I region does not readily aggregate when expressed alone. (**F**) The S region does not readily aggregate when expressed alone. (**G**) The C region does not readily aggregate when expressed alone. All DISC1 constructs shown here are Flag-tagged and visualised using an anti-Flag antibody. Western blots are from HEK293 cell lysates. Microscopy images are from SH-SY5Y cells, and are representative of three independent experiments, with scale bars representing 10 μm.

Therefore, we next investigated combinations of these vectors, to see whether these instead led to aggregation (figure 3A). Notable, while neither the I and S regions paired together (AA 539-738, figure 3B), nor the S and C regions paired together formed aggregates (AA 539-738, figure 3C), the combination of all three of the I, S and C regions did lead to consistent aggregation (AA 539-854, figure 3D). Separately, a construct encoding both the D and I regions, along with the uncharacterised “linker” region between them demonstrated consistent aggregation (AA 257-655, figure 3E). This region also formed aggregates if it was terminated in the middle of the I region, at the site of a chromosomal translocation breakpoint, found in a family with unusually high loading of mental illness [13,38,39] (AA 257-598, figure 3F). It therefore appears that DISC1 can aggregate through at least two distinct mechanisms, one based on central regions of the protein, and one based on C-terminal parts of it.

**Figure 3:**
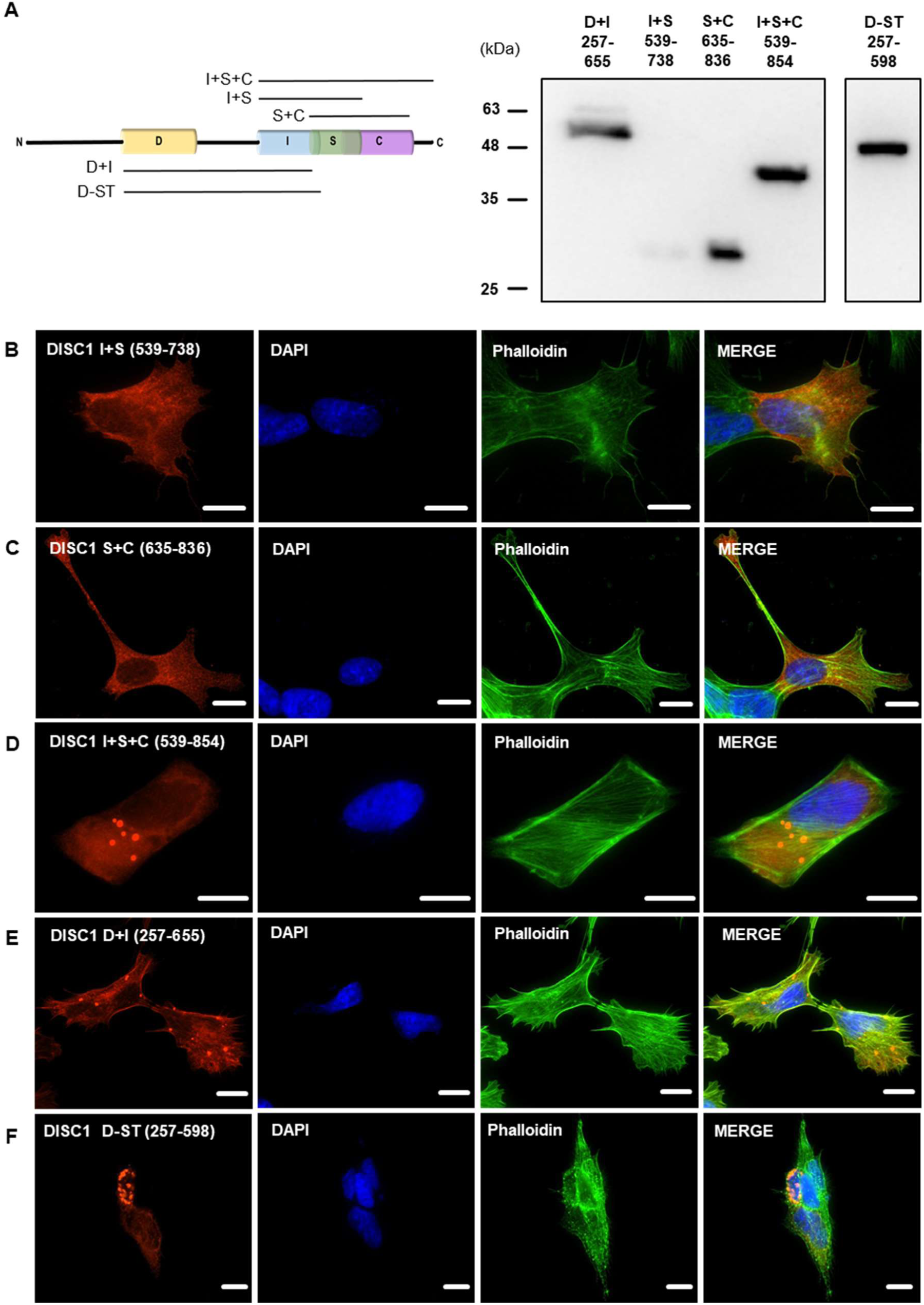
Some combinations of DISC1 regions form aggregates. (**A**) Constructs encoding multiple regions of DISC1, shown in schematic form and by Western blot. (**B**) A DISC1 construct consisting of the I and S regions do not readily aggregate when expressed. (**C**) A DISC1 construct consisting of the S and C regions do not readily aggregate when expressed. (**D**) A DISC1 construct consisting of the I, S and C regions consistently forms aggregates when expressed. (**E**) A DISC1 construct consisting of the D and I regions, plus the linker between them, consistently forms aggregates when expressed. (**D**) A DISC1 construct consisting of the D region and extending to a known chromosomal translocation breakpoint in the middle of the I region, consistently forms aggregates when expressed. All DISC1 constructs shown here are Flag-tagged and visualised using an anti-Flag antibody. Western blots are from HEK293 cell lysates. Microscopy images are from SH-SY5Y cells, and are representative of three independent experiments, with scale bars representing 10 μm.

We then generated constructs to investigate this potential aggregation site near the centre of the protein in more detail (figure 4A). When only the D region and this linker were expressed, aggregates were once again seen (AA 257-538, figure 4B), a result that was also seen using a construct expressing the linker region and the putative I region (AA 384-655, figure 4C). This strongly implicates the linker region itself as the cause of aggregation, which was confirmed by expressing it alone (AA 384-538, figure 4D). To confirm this, we performed an insolubility assay, in which HEK293 cells were transfected with a plasmid encoding either the D region alone, the D region with linker or the linker alone. These were then lysed, and had their insoluble protein fractions purified through a number of ultracentrifugation steps in different buffers. All three fragments were present in the original cell lysate, but notably only the constructs containing the linker region alone, or the linker region plus D, were seen in the purified insoluble fraction, while the D region alone was not (figure 4E). Given that protein aggregates normally show a much higher degree of insolubility than the equivalent proteins in a correctly folded state [1], this supports the idea that the linker region is responsible and sufficient for aggregation of DISC1.

**Figure 4:**
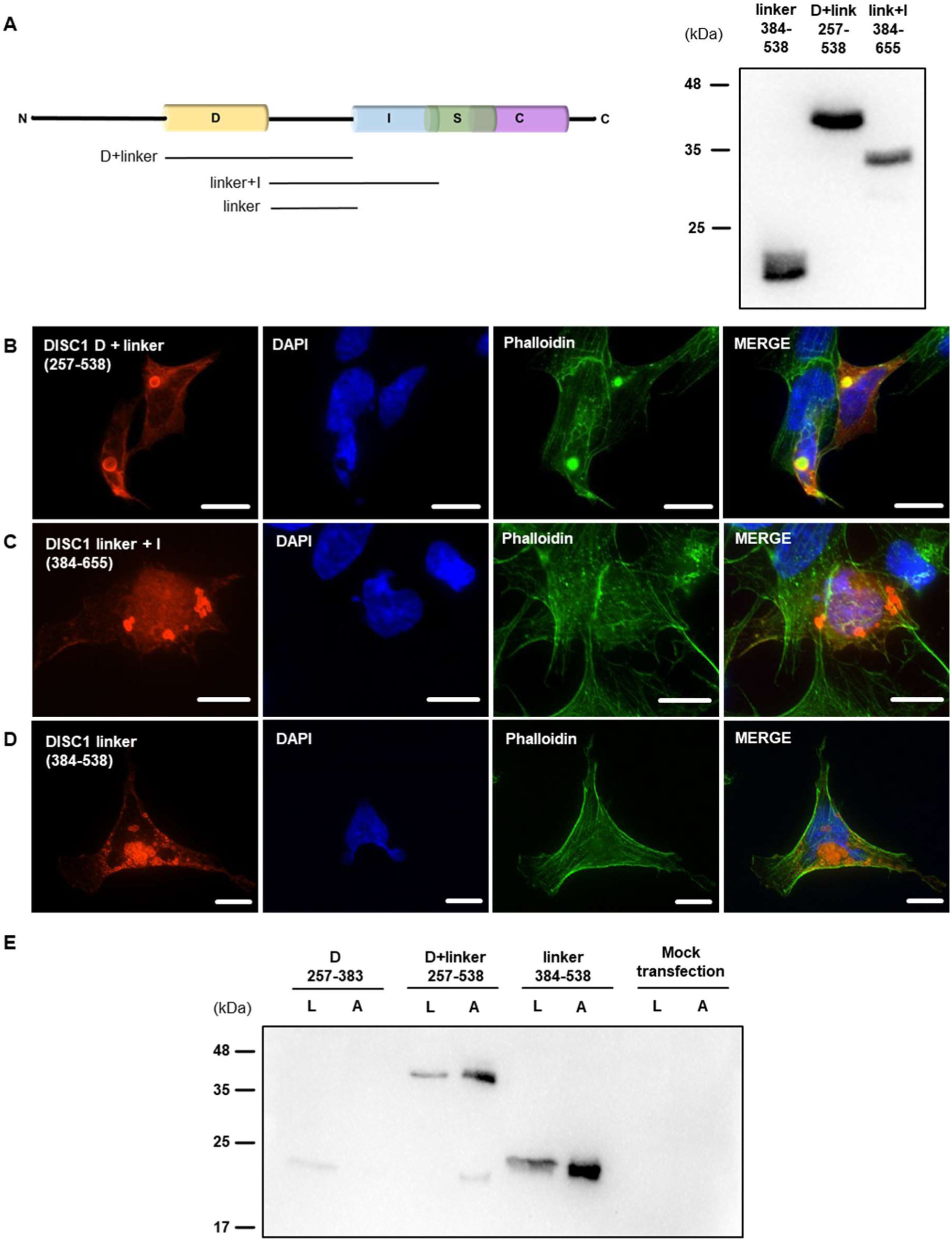
The linker region is sufficient to induce DISC1 aggregation. (**A**) Constructs encoding the linker region with or without the D or I regions, shown in schematic form and by Western blot. (**B**) A DISC1 construct consisting of the D region and linker region consistently forms aggregates when expressed. (**C**) A DISC1 construct consisting of the linker region and I region consistently forms aggregates when expressed. (**D**) A DISC1 construct consisting of the linker region alone consistently forms aggregates when expressed. (**E**) An insolubility assay, in which cells are transfected with DISC1 constructs, lysed, and then have their insoluble protein faction purified through a series of solubilisation and ultracentrifugation steps. All DISC1 fragments are visible in the crude cell lysates (L), while only those containing the linker region are enriched in the purified insoluble fraction, where aggregates are expected to be found (A). All DISC1 constructs shown here are Flag-tagged and visualised using an anti-Flag antibody. Western blots are from HEK293 cell lysates. Microscopy images are from SH-SY5Y cells, and are representative of three independent experiments, with scale bars representing 10 μm.

To further confirm this, constructs were generated that encoded DISC1 from its N-terminus through to the D region or part way through the linker region (figure 5A). Strikingly, while a construct ending at the classic C-terminal end of the D region did not form aggregates (AA 1-383, figure 5B), one ending 32 amino acids later did (AA 1-415, figure 5C), indicating that this region is critical for aggregation. Finally, to confirm this, a full-length version of DISC1 was generated in which these 32 amino acids were deleted. DISC1 constructs lacking this region consistently failed to aggregate (AA1-854 Δ384-414, AA 1-836 Δ384-414 and AA 1-655 Δ384-414, 5D-F), while full length DISC1 did form aggregates under the same circumstances (figure 5G).

**Figure 5:**
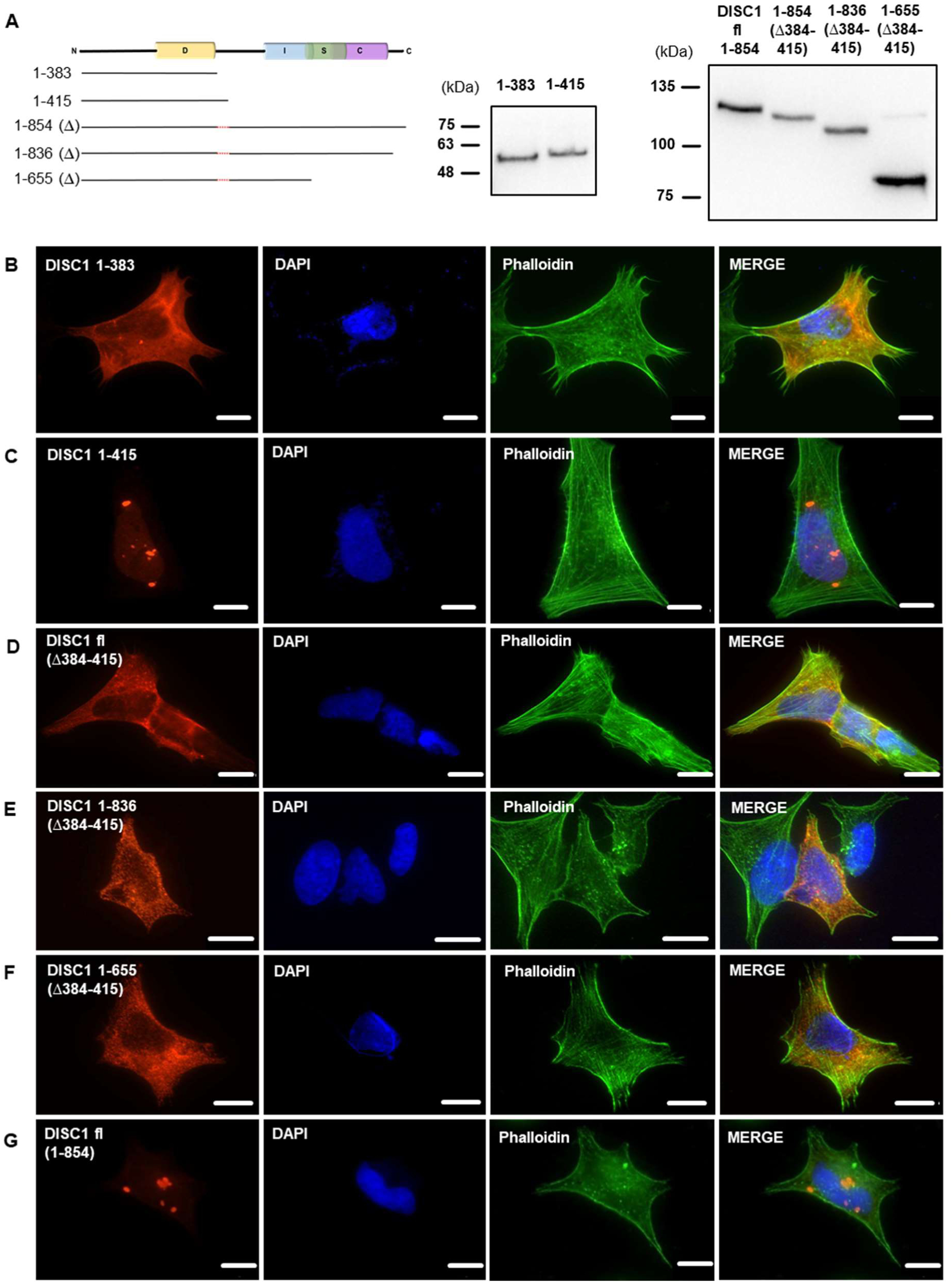
Deletion of part of the linker region abolishes the aggregation propensity of DISC1. (**A**) Constructs encoding DISC1 or fragments of it, shown in schematic form and by Western blot. (**B**) A DISC1 construct consisting of the N-terminus of DISC1 up to the beginning of the linker region does not readily aggregate when expressed. (**C**) A DISC1 construct consisting of the N-terminus of DISC1 up to part way through the linker region forms aggregates when expressed. (**D**) Full length DISC1 missing this section of the linker region does not readily aggregate. (**E**) Full length DISC1 missing this section of the linker region and sequence after the C region does not readily aggregate. (**F**) Full length DISC1 missing this section of the linker region and sequence after the I region does not readily aggregate. (**G**) Full length DISC1 consistently forms aggregates. All DISC1 constructs shown here are Flag-tagged and visualised using an anti-Flag antibody. Western blots are from HEK293 cell lysates. Microscopy images are from SH-SY5Y cells, and are representative of three independent experiments, with scale bars representing 10 μm.

## Discussion

DISC1 has previously been determined to aggregate in the brains of patients with schizophrenia, bipolar disorder and depression [5,25], with expression of aggregating DISC1 in the rat model leading to a variety of neuroanatomical, behavioural and dopaminergic changes, reminiscent of mental illness [18,19,22,42,43]. Here, we investigated whether a specific region of DISC1 was required for its aggregation, determining that AA 384-415, immediately C-terminal of the D region, was both sufficient for aggregation, and that aggregation was abolished when it was deleted. The exact structure of this region is unknown. It is not found within any of 94 most stable DISC1 fragments yielded by the ESPRIT screening, but would be predicted to be α-helical based on secondary structure prediction [28]. It remains unclear, however, whether aggregation of DISC1 in the cellular environment occurs through the same mechanism as the propensity of purified recombinant DISC1 to aggregate. It is also notable that the insoluble (aggregated) DISC1 detected in the brains of patients is not the 100kDa full length protein, but a species of around 70 kDa, which was detected using an antibody that recognises part of the C region of DISC1 [5,32]. The exact nature of this species is unclear, however if it is the result of posttranslational processing, then based on its size and the fact that the C-terminus is present, it would be expected to have lost approximately the first 250 amino acids. While it is not possible to accurately determine the processing site from existing data, this is consistent with a species that begins with the D region, and extends to the C-terminus of the full-length protein, thus containing all known structured regions and the aggregation-critical region described here. To our knowledge, however, no one has attempted to study insoluble DISC1 in human brain tissue using an antibody against the N-terminus of the protein. Our results are somewhat similar to an early DISC1 study that looked at “punctate staining” of protein fragments expressed in HeLa cells, which also saw central regions of DISC1 to be important, but which concluded a location in what is now considered the I region to be responsible [44]. The differences between their results and ours in SH-SY5Y may arise because their construct boundaries were defined prior to an understanding of the DISC1 domain structure, meaning that their constructs represented partial domains that were unstable as a result.

A previous study has also implicated a short region of DISC1, N-terminal of the D region (AA 209-227) as being critical for aggregation with HTT, based on purified protein [45]. Specifically, this region is critical for interaction of DISC1 with HTT, as determined by a peptide array, and it appears that HTT can induce aggregation of previously stable DISC1 [45]. That we found a separate region of DISC1 to be required for its aggregation in cells, suggests that auto-aggregation of DISC1 likely occurs through distinct mechanisms than those by which HTT can induce DISC1 to co-aggregate. This is also similar to the way in which DISC1 can induce TRIOBP-1 to co-aggregate with it, even if the regions of TRIOBP-1 known to be responsible for its auto-aggregation are deleted [11,12].

Another study noticed that a C-terminal region of DICS1, AA 640-854, approximating to the S and C regions, was capable of aggregation in cells based on an insolubility assay, and also that insoluble C-terminal fragments of DISC1 were found in patient’s brains [25]. These results broadly align with our results that there is a distinct, as yet undetermined, mechanism by which the I, S and C regions (AA 539-854) could form aggregates, although we saw no such propensity when the S and C regions (AA 635-836) were expressed instead. It also cannot be formally ruled out that the C-terminal 18 amino acids of DISC1 may play a role in aggregation. More recently, a biophysical characterisation of the C region showed it to be capable of aggregating *in vitro*, with these aggregates seen to take the form of β-fibrils [40]. This self-association and ultimately aggregation of the C region was driven on a “β-core” region from AA 716-761, and impacted the binding of DISC1 to key interaction partners [40].

These two different aggregation critical regions, the linker between the D and I regions and in the C region, are not necessarily incompatible, given that both were determined in different systems. In neuroblastoma cells, the region between D and I shows the highest aggregation capacity of any individual section of DISC1 examined, dwarfing any effect of aggregation of the C region alone. When using recombinant protein, the C region is the easiest section of DISC1 to express and purify from *E. coli* [32], but can be induced to aggregate [40], while in comparison, the linker region between D and I is sufficiently unstable that it is very hard to express and purify under the same circumstances (unpublished observation by us, Abhishek Cukkemane and Oliver Weiergräber). It is therefore likely that the linker region has the inherently higher aggregation propensity, when studied in isolation, making it dominant in our neuroblastoma system, and making it impractical to work with as a recombinant protein. In contrast, the subtler aggregation of the C region can be studied using recombinant proteins, but is largely masked in neuroblastoma cells (excepting, perhaps, when co-expressed with the I and S regions). This does not necessarily mean that the linker region is more relevant to pathological aggregation than the C region, however, given that both regions were determined using model systems, not endogenous protein, and given that other cell-based studies, with lower levels of over-expressed DISC1, showed DISC1 to aggregate only after induction, in that case by dopamine [18].

It therefore appears that DISC1 aggregation, which is implicated in schizophrenia, bipolar disorder and major depressive disorder [5], may arise through more than one molecular mechanism. It was previously known that aggregation could arise, at least in recombinant protein, through the C region [40], and we now demonstrate that in mammalian cell systems, aggregation is dependent on a short linker region between the D and I regions. This information, and in particular the generation of a slightly truncated form of DISC1 lacking its aggregation propensity, can now be used to generate better model systems for studying DISC1 aggregation, through separating out effects caused by over-expression of DISC1, from those caused specifically by its aggregation.

## Acknowledgements

This work was funded by grants from the Croatian Science Foundation (IP-2018-01-9424, DOK-2018-09-5395) and the Alexander von Humboldt Foundation (1142747-HRV-IP). We thank Abhishek Cukkemane and Oliver Weiergräber for their guidance and advice on recombinant protein expression, Bobana Samardžija for experimental assistance, and Elizabeth Bradshaw for proofreading.

## Author contributions

BZ performed the experiments. Both authors designed the experiments and analysed the results. NJB drafted the paper and BZ prepared figures. Both authors edited and approved the final version.

