## Supplementary tables S1 & S2 for "Aggregation of Disrupted in Schizophrenia 1 arises from a central region of the protein"

### Supplementary material

| | |
|---|---|
| Table S1 | Details of plasmids used in this study |
| Table S2 | Details of primers used in this study |

**Table S1**: Details of all plasmid vectors used for cloning in this study. See table S2 for details of each primer used in generating them

| No. | Vector backbone | Gene insert | Origin |
|---|---|---|---|
| 1 | pENTR1A no ccDB | (None) | Addgene, clone 17398, Campeau et al (2009) PLOS One 4:e6529 |
| 2 | pDONR/Zeo | (None) | Thermo Fisher Scientific |
| 3 | pdcDNA-FlagMyc | (None) | BCCM/LMBP Plasmid Collection, clone LMBP 4705 |
| 4 | pETG10A | (None) | A. Geerlof, EMBL, Heidelberg, Germany |
| 5 | pENTR223 | DISC1 (full length) | DNASU Plasmid Repository, clone HsCD00516321 |
| 6 | pENTR1A | DISC1 (1-383, Nterm-D) | Subcloned from vector 5 in two rounds using primers A and J, then J and T, before being ligated into SalI/KpnI sites of vector 1 |
| 7 | pENTR1A | DISC1 (1-415, Nterm-linker) | Subcloned from vector 5 in two rounds using primers A and L, then L and T, before being ligated into SalI/KpnI sites of vector 1 |
| 8 | pDONR/Zeo | DISC1 (257-383, D) | Subcloned from vector 5 in two rounds using primers C and J, then J and V, before being transferred into vector 2 using BP clonase |
| 9 | pENTR1A | DISC1 (257-538, D-linker) | Subcloned from vector 5 in two rounds using primers B and M, then T and U, before being ligated into SalI/XbaI sites of vector 1 |
| 10 | pENTR1A | DISC1 (257-597, D-I) | Subcloned from vector 5 in two rounds using primers B and N, then T and U, before being ligated into SalI/XbaI sites of vector 1 |
| 11 | pENTR1A | DISC1 (257-655, D-I) | Subcloned from vector 5 in two rounds using primers B and O, then T and U, before being ligated into SalI/XbaI sites of vector 1 |
| 12 | pENTR1A | DISC1 (384-538, linker) | Subcloned from vector 5 in two rounds using primers D and M, then T and U, before being ligated into SalI/XbaI sites of vector 1 |
| 13 | pENTR1A | DISC1 (384-655, linker-I) | Subcloned from vector 5 in two rounds using primers D and O, then T and U, before being ligated into SalI/XbaI sites of vector 1 |
| 14 | pENTR1A | DISC1 (539-655, I) | Subcloned from vector 5 in two rounds using primers F and O, then T and U, before being ligated into SalI/XbaI sites of vector 1 |
| 15 | pENTR1A | DISC1 (539-738, I-S) | Subcloned from vector 5 in two rounds using primers F and O, then T and U, before being ligated into SalI/XbaI sites of vector 1 |
| 16 | pENTR1A | DISC1 (539-854, I-C) | Subcloned from vector 5 in two rounds using primers F and S, then T and U, before being ligated into SalI/XbaI sites of vector 1 |
| 17 | pDONR/Zeo | DISC1 (635-738, S) | Subcloned from vector 5 in two rounds using primers H and Q, then Q and V, before being transferred into vector 2 using BP clonase |

| No. | Vector backbone | Gene insert | Origin |
|---|---|---|---|
| 18 | pENTR1A | DISC1 (635-836, S-C) | Subcloned from vector 5 in two rounds using primers G and R, then T and U, before being ligated into SalI/XbaI sites of vector 1 |
| 19 | pENTR1A | DISC1 (691-836, C) | Subcloned from vector 5 in two rounds using primers I and R, then T and U, before being ligated into SalI/XbaI sites of vector 1 |
| 20 | pENTR1A | DISC1 (Δ384-414) | Subcloned from vector 5 in two rounds using primers E and S, then E and V, before being ligated into KpnI/XbaI sites of vector 5 |
| 21 | pENTR1A | DISC1 (1-655Δ384-414) | Subcloned from vector 20 in two rounds using primers O and T, then T and U, before being ligated into SalI/XbaI sites of vector 1 |
| 22 | pENTR1A | DISC1 (1-836Δ384-414) | Subcloned from vector 20 in two rounds using primers R and T, then T and U, before being ligated into SalI/XbaI sites of vector 1 |
| 23 | pdcDNA-Flag | DISC1 (full length) | Samardžija, Juković et al (2023) Cells 12:1848 |
| 24 | pdcDNA-Flag | DISC1 (1-383, Nterm-D) | LR clonase recombination of vectors 3 and 6 |
| 25 | pdcDNA-Flag | DISC1 (1-415, Nterm-linker) | LR clonase recombination of vectors 3 and 7 |
| 26 | pdcDNA-Flag | DISC1 (257-383, D) | LR clonase recombination of vectors 3 and 8 |
| 27 | pdcDNA-Flag | DISC1 (257-538, D-linker) | LR clonase recombination of vectors 3 and 9 |
| 28 | pdcDNA-Flag | DISC1 (257-597, D-I) | LR clonase recombination of vectors 3 and 10 |
| 29 | pdcDNA-Flag | DISC1 (257-655, D-I) | LR clonase recombination of vectors 3 and 11 |
| 30 | pdcDNA-Flag | DISC1 (384-538, linker) | LR clonase recombination of vectors 3 and 12 |
| 31 | pdcDNA-Flag | DISC1 (384-655, linker-I) | LR clonase recombination of vectors 3 and 13 |
| 32 | pdcDNA-Flag | DISC1 (539-655, I) | LR clonase recombination of vectors 3 and 14 |
| 33 | pdcDNA-Flag | DISC1 (539-738, I-S) | LR clonase recombination of vectors 3 and 15 |
| 34 | pdcDNA-Flag | DISC1 (539-854, I-C) | LR clonase recombination of vectors 3 and 16 |
| 35 | pdcDNA-Flag | DISC1 (635-738, S) | LR clonase recombination of vectors 3 and 17 |
| 35 | pdcDNA-Flag | DISC1 (635-836, S-C) | LR clonase recombination of vectors 3 and 18 |
| 36 | pdcDNA-Flag | DISC1 (691-836, C) | LR clonase recombination of vectors 3 and 19 |
| 37 | pdcDNA-Flag | DISC1 (Δ384-414) | LR clonase recombination of vectors 3 and 21 |
| 38 | pdcDNA-Flag | DISC1 (1-655Δ384-414) | LR clonase recombination of vectors 3 and 22 |
| 39 | pdcDNA-Flag | DISC1 (1-836Δ384-414) | LR clonase recombination of vectors 3 and 20 |
| 40 | pETG10A | DISC1 (384-538, linker) | LR clonase recombination of vectors 4 and 12 |

**Table S2**: Details of all primers used for cloning in this study.

| ID | Name | Sequence | Purpose |
|---|---|---|---|
| A | DISC1-1-salF | GTAGTCGACATGCCAGGCGG | Forward primer for cloning DISC1 from amino acid 1. Adds a SalI site. |
| B | DISC1-257-salF | GTAAGTCGACATGGAGGACCCGC | Forward primer for cloning DISC1 from amino acid 257. Adds a SalI site. |
| C | DISC1-257-attbF | AATAGCAGGCTTCGCCGCCACCATGGAGGACCCGCG | Forward primer for cloning DISC1 from amino acid 257. Adds part of an attB1 recombination site. |
| D | DISC1-384-salF | GGCGTCGACATGGAACAAGAGAAAATC | Forward primer for cloning DISC1 from amino acid 384. Adds a SalI site. |
| E | DISC1-415-salkpnF | GAAAAGTCGACATGGGTACCGCCTTGCGCC | Forward primer for cloning DISC1 from amino acid 415. Adds a SalI and a KpnI site. |
| F | DISC1-539-salF | GTAGTCGACATGCCACCGGAAAC | Forward primer for cloning DISC1 from amino acid 539. Adds a SalI site. |
| G | DISC1-635-salF | GGCGTCGACATGAATGTCAAAAAGCTG | Forward primer for cloning DISC1 from amino acid 635. Adds a SalI site. |
| H | DISC1-635-attbF | AGCAGGCTTCGCCGCCACCATGAATGTCAAAAAG | Forward primer for cloning DISC1 from amino acid 635. Adds part of an attB1 recombination site. |
| I | DISC1-691-salF | GGAGTCGACATGTGGGAAGCTG | Forward primer for cloning DISC1 from amino acid 691. Adds a SalI site. |
| J | DISC1-383-xbakpnR | GCGCGTCTAGACTATCAGGTACCCAGGTCTTCTAATC | Reverse primer for cloning DISC1 from amino acid 383. Adds an XbaI and a KpnI site. |
| K | DISC1-383-attR | GGCCACCACTTTGTACAAGAAAGCTGGGTCTCACAGGTCTTCTAATC | Reverse primer for cloning DISC1 from amino acid 383. Adds an attB2 recombination site. |
| L | DISC1-415-xbakpnR | GCAAATCTAGACTATCAGGTACCGGCAGCCTGG | Reverse primer for cloning DISC1 from amino acid 415. Adds an XbaI and a KpnI site. |
| M | DISC1-538-ecoR | GGCGAATTCTCATCACTCTGCATGG | Reverse primer for cloning DISC1 from amino acid 538. Adds an EcoRI site. |
| N | DISC1-597-ecoR | GGCGTCGACATGTCAGGAAACC | Reverse primer for cloning DISC1 from amino acid 597. Adds an EcoRI site. |
| O | DISC1-655-ecoR | GAGGAATTCTCATCAGTGCTCCACTTC | Reverse primer for cloning DISC1 from amino acid 655. Adds an EcoRI site. |

| ID | Name | Sequence | Purpose |
|---|---|---|---|
| P | DISC1-738-ecoR | GGAGAATTCTCATCAGGAGTGGAGC | Reverse primer for cloning DISC1 from amino acid 738. Adds an EcoRI site. |
| Q | DISC1-738-attbR | CACCACTTTGTACAAGAAAGCTGGGTCTCAGGAGTGGAGC | Reverse primer for cloning DISC1 from amino acid 738. Adds an attB2 recombination site. |
| R | DISC1-836-ecoR | GTAGAATTCTCATCATCCCGCCTCC | Reverse primer for cloning DISC1 from amino acid 836. Adds an EcoRI site. |
| S | DISC1-854-ecoR | GGAGAATTCTCATCAGGCTTGTGCTTC | Reverse primer for cloning DISC1 from amino acid 854. Adds an EcoRI site. |
| T | Extension F | GCTATAAGGATCCGGTACCTAGTCGACATG | Secondary forward primer, adds BamHI and KpnI sites. |
| U | Extension R | GGCACCAGCTCGAGTCTAGAATTCTCATC | Secondary reverse primes, adds XhoI and XbaI sites. |
| V | attB 5' F | GTCGACACAAGTTTGTACAAAAAAGCAGGCTTCGCCGCCACC | Secondary forward primer, completes an attB1 recombination site. |
